# Glucocorticoid stress hormones stimulate vesicle-free Tau secretion and spreading in the brain

**DOI:** 10.1101/2023.06.07.544054

**Authors:** Qing Yu, Fang Du, Irla Belli, Patricia A. Gomes, Ioannis Sotiropoulos, Clarissa L. Waites

## Abstract

Chronic stress and elevated levels of glucocorticoids (GCs), the main stress hormones, accelerate Alzheimer’s disease (AD) onset and progression. A major driver of AD progression is the spreading of pathogenic Tau protein between brain regions, precipitated by neuronal Tau secretion. While stress and high GC levels are known to induce intraneuronal Tau pathology (*i.e.* hyperphosphorylation, oligomerization) in animal models, their role in trans-neuronal Tau spreading is unexplored. Here, we find that GCs promote secretion of full-length, vesicle-free, phosphorylated Tau from murine hippocampal neurons and *ex vivo* brain slices. This process occurs via type 1 unconventional protein secretion (UPS) and requires neuronal activity and the kinase GSK3β. GCs also dramatically enhance trans-neuronal Tau spreading *in vivo*, and this effect is blocked by an inhibitor of Tau oligomerization and type 1 UPS. These findings uncover a potential mechanism by which stress/GCs stimulate Tau propagation in AD.

## Introduction

Stressful life events and high circulating levels of glucocorticoids (GCs), the primary stress hormones, are known risk factors for Alzheimer’s disease (AD)^1–3^. Indeed, epidemiological and clinical studies suggest that prolonged psychosocial stress significantly elevates AD risk, and high GC levels are associated with faster cognitive decline in AD patients^1–5^. Moreover, stress interacts with genetic risk factors to hasten the onset of AD symptoms and pathology in both animal models and humans^6–12^. Stress and AD appear to share a common pathological driver: the microtubule-associated protein Tau. Not only do stress and high GC levels trigger Tau pathology similar to that seen in AD brain tissue (*i.e*., Tau hyperphosphorylation and aggregation)^13–15^, but Tau depletion is protective against both amyloid beta-and stress-induced neurotoxicity and cognitive impairment in animal models^15–17^, indicating Tau’s essential role as a mediator of neurodegeneration in the context of AD and chronic stress.

A key feature of Tau pathology in AD is its stereotypical spreading pattern between anatomically connected brain regions (entorhinal cortex to hippocampus to prefrontal cortex)^18^. This spreading is highly correlated with the severity of cognitive impairment in AD patients and appears to be a major driver of AD progression^18–20^. Given the relationship between Tau propagation and clinical AD symptoms, there has been tremendous interest in elucidating the mechanisms of Tau secretion and spreading in the brain. Numerous studies have shown that Tau is secreted from neurons in extracellular vesicles, including ectosomes that derive from the plasma membrane and exosomes that derive from multivesicular endosomes of the endolysosomal pathway, and also as vesicle-free protein^21–23^. While vesicle-mediated mechanisms of Tau spreading have been a focus of study for over a decade^21, 24, 25^, the vast majority of Tau secreted by neurons (∼90%) is vesicle-free^23, 25–32^, and considerably less is known about this mode of secretion and its contribution to pathogenic Tau propagation. Similarly, although chronic stress and high GC levels are known to induce Tau pathology in the hippocampus and cortex, precipitating synaptic loss and behavioral impairment in animal models (*i.e.* anxiety, anhedonia, learning/memory deficits)^14, 15, 33^, it is unclear whether or how stress/GCs stimulate the spreading of Tau pathology between these brain regions.

In the current study, we investigate the effects of GCs on neuronal Tau secretion and spreading in murine hippocampal neurons, brain slices, and *in vivo* hippocampus. We find that GCs induce secretion of vesicle-free Tau through type 1 unconventional protein secretion (UPS), in an activity-and glycogen synthase kinase 3β (GSK3β)-dependent manner. Moreover, GC administration stimulates Tau spreading through the hippocampus, and this process is prevented by inhibiting Tau aggregation and type 1 UPS with the catechin EGCG. Together, these findings demonstrate that elevated GC levels promote Tau propagation, and suggest a mechanism by which stress/GCs speeds cognitive decline in AD.

## Results

To determine whether GCs stimulate neuronal Tau secretion, we measured extracellular Tau levels by immunoblot and/or ELISA in three preparations: 1) media from 14 day *in vitro* (DIV) murine hippocampal neurons treated for 48 hours with vehicle control, the synthetic GC dexamethasone (DEX), or DEX + GC receptor (GR) antagonist mifepristone (MIF)(Fig. **1A-D, I**), 2) artificial cerebrospinal fluid (ACSF) from *ex vivo* murine brain slices of 4 months old mice, perfused for 4 hours with vehicle, DEX, or DEX + MIF (Fig. **1E-H, J**), and 3) CSF from 4-5 months old mice administered vehicle, DEX, or DEX + MIF for 15 days (Fig. **1K**). The efficacy of DEX treatment was confirmed by immunoblotting hippocampal lysates for phospho-GR and by immunostaining for phospho-and oligomeric Tau (Fig. **S1A-E**), as in our recent study^34^. In all three preparations, DEX significantly increased Tau concentration compared to vehicle and DEX + MIF (Fig. **1A-K**). This increase in extracellular Tau did not result from cell death or disruption of plasma membrane integrity, as lactate dehydrogenase (LDH) levels were unaltered by DEX +/- MIF in both the *in vitro* and *ex vivo* preparations (Fig. **1L**), and the abundant cytoskeletal proteins actin and tubulin were not detected in these fluids (Fig. **1A, E**). We also found that extracellular Tau was predominantly full-length, phosphorylated at multiple sites, as indicated by immunoreactivity for AT8 (Ser202/Thr205) and PHF1 (Ser396/Ser404) antibodies (Fig. **1A-C, E-G**), and vesicle-free rather than associated with extracellular vesicles (EVs), which were depleted from media and ACSF by a well-established centrifugation procedure (Fig. **S1F-H**)^35, 36^. Extracellular Tau levels were similarly increased in media containing cortical and hippocampal brain slices from mice subjected chronic unpredictable stress (CUS) compared to control conditions (Fig. **S1I**), demonstrating that CUS and GC exposure have comparable stimulatory effects on Tau secretion.

**Figure 1.**
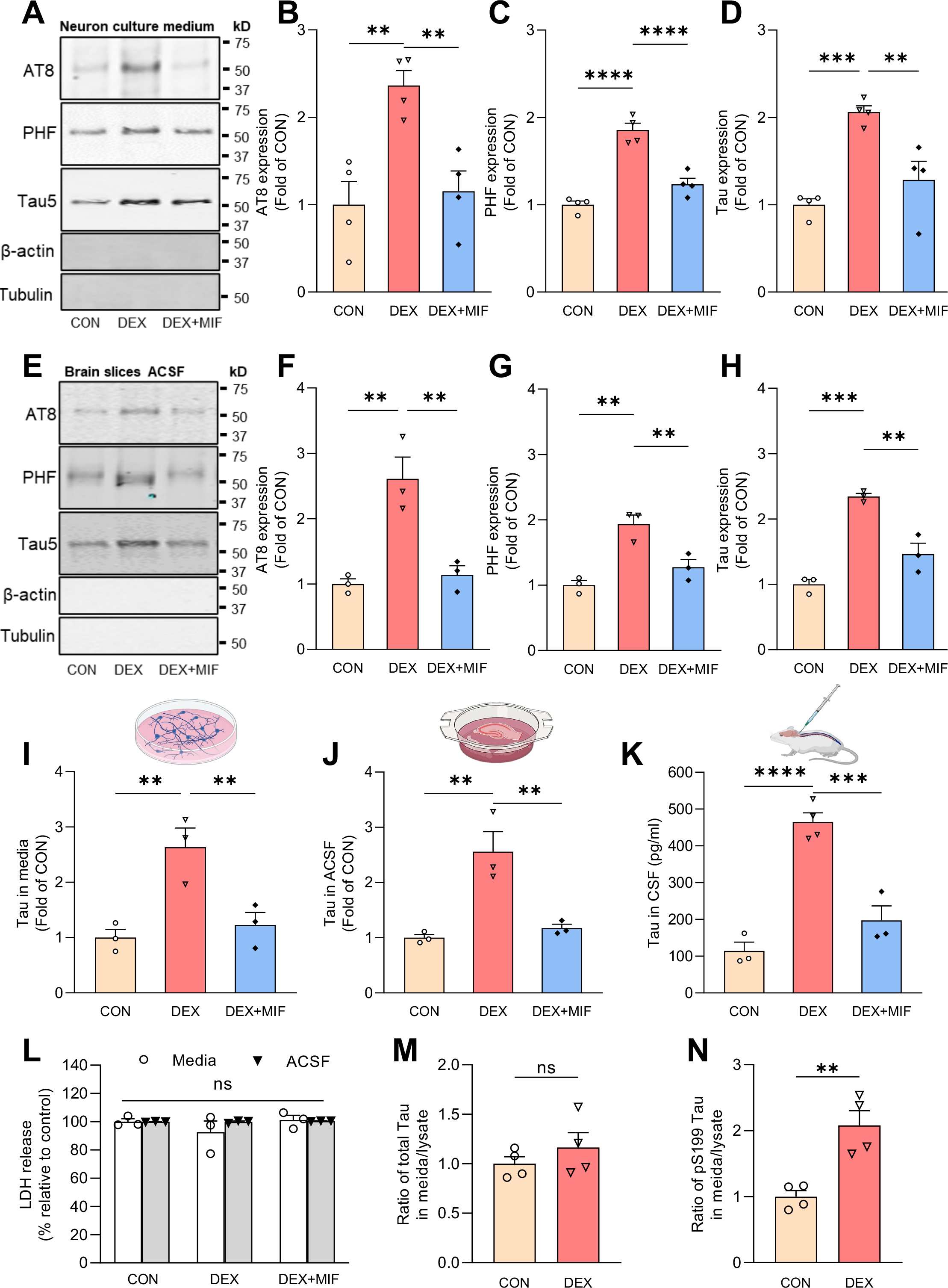
Glucocorticoids induce secretion of vesicle-free, phosphorylated Tau. (**A-D**) Representative immunoblots (**A**) and quantification **(B-D)** of AT8, PHF1, and total Tau (Tau5) immunoreactivity in extracellular vesicle (EV)-depleted media from hippocampal neurons treated with vehicle (CON), DEX, or DEX+MIF. Intensity values are normalized to the CON condition (**P _CON VS. DEX_ =0.0021, **P_DEX VS. DEX + MIF_ = 0.0043 for **B**, ****P_CON VS. DEX_ <0.0001, ****P_DEX VS. DEX + MIF_ <0.0001 for **C**, ***P_CON VS. DEX_=0.0004, **P_DEX VS. DEX + MIF_=0.0029 for **D**, one-way ANOVA with multiple comparisons and Fisher’s LSD test; n=4 samples/condition). (**E-H**) Representative immunoblots (**E**) and quantification **(F-H)** of AT8, PHF1, and total Tau (Tau5) immunoreactivity in EV-depleted ACSF from brain slices perfused with vehicle (CON), DEX, or DEX + MIF. Intensity values are normalized to the CON condition (**P _CON VS. DEX_ =0.0017, **P_DEX VS. DEX + MIF_ =0.0027 for **F**, **P_CON VS. DEX_ =0.0011, **P_DEX VS. DEX + MIF_ =0.0061 for **G**, ***P_CON VS. DEX_=0.0001, **P_DEX VS. DEX + MIF_ =0.0012 for **H**, one-way ANOVA with multiple comparisons and Fisher’s LSD test; n=3 samples/condition). (**I**-**K)** Quantification of ELISA for total Tau levels in EV-depleted media from hippocampal neurons (**I**), ACSF from brain slices (**J**), and CSF from mice (**K**) following the indicated treatments. Values are normalized to CON condition in **I**-**J** and expressed as pg/ml in **K** (**P _CON VS. DEX_ =0.0039, **P_DEX VS. DEX + MIF_ =0.0079 for **I**, **P_CON VS. DEX_ =0.0023, **P_DEX VS. DEX + MIF_ =0.0041 for **J**, ****P_CON VS. DEX_<0.0001, ***P_DEX VS. DEX + MIF_ =0.0003 for **K**, one-way ANOVA with multiple comparisons and Fisher’s LSD test; n=3 samples/condition). (**L**) Quantification of LDH in extracellular vesicle (EV)-depleted media (white bars) or ACSF (gray bars) from the indicated treatment conditions (*P*_CON VS. DEX_ =0.1718, *P*_DEX VS. DEX + MIF_ =0.1185 for media samples, *P*_CON VS. DEX_ =0.9905, *P*_DEX VS. DEX + MIF_ =0.9191 for ACSF samples, two-way ANOVA with multiple comparisons and Fisher’s LSD test; *n*=3 samples/condition). (**M-N**) Ratio of Tau concentration in neuronal media to Tau concentration in neuronal lysate for CON and DEX conditions, measured by ELISA, for total Tau (**M**) and pS199 Tau (**N**)(**P_CON VS. DEX_ =0.0041, unpaired two-tailed t-test, n=4 samples/condition).

Since stress/GCs promote Tau accumulation, these findings could reflect similar levels of Tau secretion from a larger intraneuronal pool. To determine whether GCs alter the fractional amount of Tau secreted from neurons, we measured Tau concentration in media versus neuronal lysate for control and DEX conditions, using ELISA kits to detect total or pS199 phospho-Tau. Interestingly, DEX treatment did not change the secreted versus intracellular ratio for total Tau (Fig. **1M**), but significantly increased this ratio for pS199 Tau (by two-fold; Fig. **1N**). These results indicate that DEX preferentially stimulates secretion of this phospho-Tau species.

Secretion of vesicle-free Tau has been shown to occur through type 1 UPS, wherein Tau is directly translocated across the plasma membrane through interactions with heparin sulfate proteoglycans (HSPGs), cholesterol, and sphingolipids^37, 38^. To determine whether GCs stimulate Tau secretion via this pathway, we first treated hippocampal neurons with vehicle or DEX +/- NaClO_3_, an inhibitor of HSPG synthesis previously shown to decrease Tau secretion via type 1 UPS^37, 38^. Following media collection, EV depletion, and measurement of phospho-and total Tau levels by immunoblot and ELISA, respectively, we found that NaClO_3_ almost completely blocked the DEX-induced increase in extracellular Tau levels (Fig. **2A-D**). Treatment with methyl-β- cyclodextrin to extract membrane cholesterol similarly inhibited DEX-induced Tau secretion (Fig. **2E-H**), demonstrating its HSPG-and cholesterol-dependence. Comparable results were seen in brain slices treated with NaClO_3_ and methyl-β-cyclodextrin (Fig. **S2A-H**). Since type 1 UPS is ATP-independent, we tried to confirm this aspect of DEX-induced Tau secretion. Unfortunately, the different time courses of DEX treatment vs. ATP depletion with 2-deoxyglucose, and the toxicity of this latter treatment, prevented us from testing both conditions simultaneously. However, we verified that baseline Tau secretion in our neuronal cultures was ATP-independent, by briefly incubating hippocampal neurons from PS19 mice (overexpressing human P301S mutant Tau; hTau) with 2-deoxyglucose (30 mM, 1 hr). While this treatment reduced cellular ATP production by ∼70%, it did not change the concentration of extracellular Tau in EV-depleted medium (Fig. **S2I**), confirming the overall ATP-independence of Tau secretion measured in our assays.

**Figure 2.**
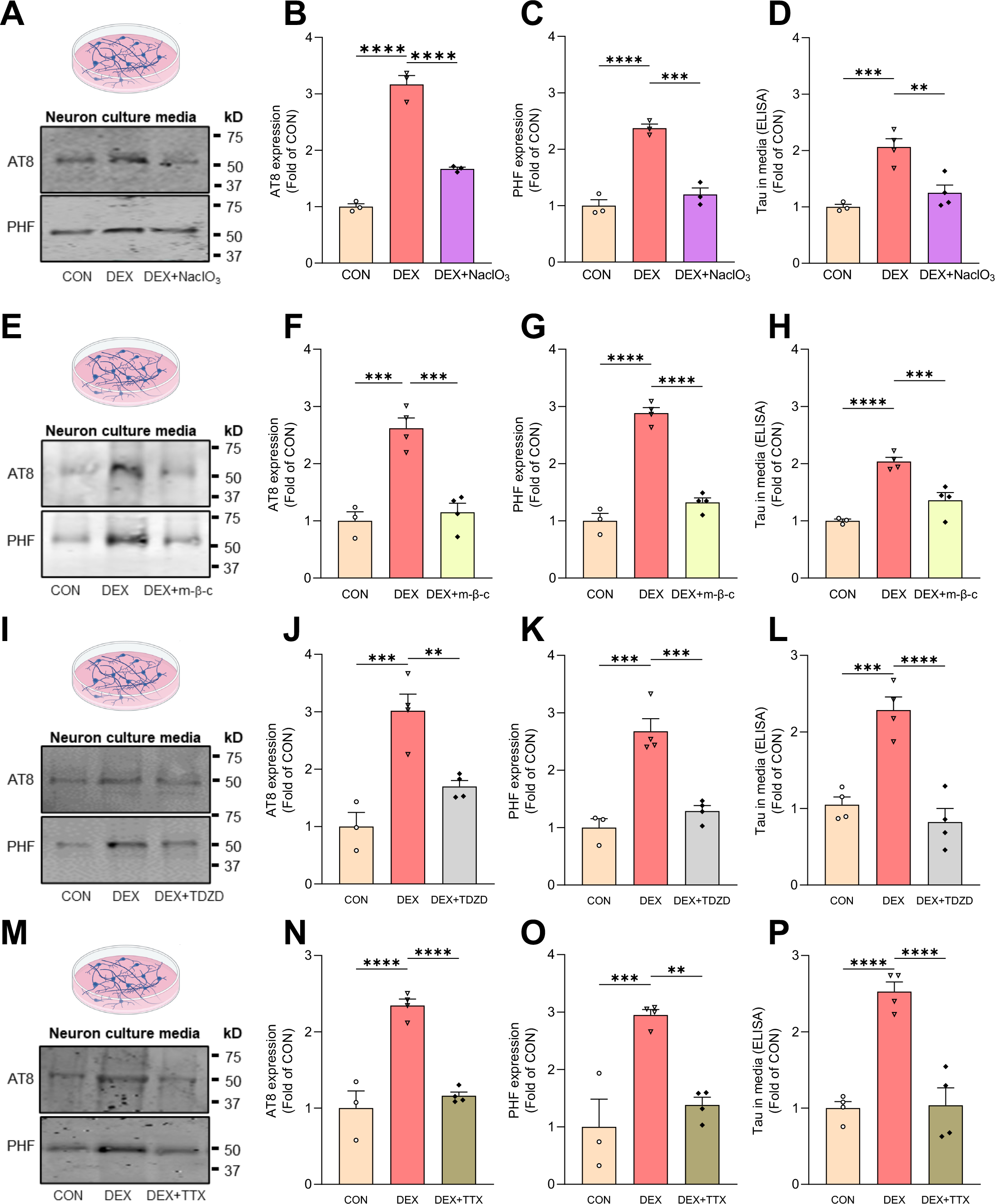
GC-mediated Tau secretion occurs via type 1 unconventional protein secretion. (**A-C**) Representative immunoblots (**A**) and quantification **(B-C)** of AT8 and PHF1 immunoreactivity in media from hippocampal neurons treated with vehicle (CON), DEX, or DEX + NaClO_3_. Intensity values are normalized to CON condition (****P _CON VS. DEX_ <0.0001, ****P_DEX VS. DEX + NaClO3_ <0.0001for **B**, ****P_CON VS. DEX_ <0.0001, ***P_DEX VS. DEX + NaClO3_ =0.0002 for **C**, one-way ANOVA with multiple comparisons and Fisher’s LSD test; n=3 samples/condition). **(D)** Quantification of ELISA for total Tau levels in EV-depleted media from the indicated conditions, with values normalized to CON condition (***P_CON VS. DEX_=0.0005, **P_DEX VS. DEX + NaClO3_ =0.0017, one-way ANOVA with multiple comparisons and Fisher’s LSD test; n=3 samples/condition). (**E-G**) Representative immunoblots (**E**) and quantification **(F-G)** of AT8 and PHF1, immunoreactivity in media from hippocampal neurons treated with vehicle (CON), DEX, or DEX + methyl-β- cyclodextrin (m-β-c). Intensity values are normalized to CON condition (***P _CON VS. DEX_ =0.0002, ***P_DEX VS. DEX + m-β-c_ =0.0002 for **F**, ****P_CON VS. DEX_ <0.0001, ****P_DEX VS. DEX + m-β-c_ <0.0001 for **G**, one-way ANOVA with multiple comparisons and Fisher’s LSD test; n=4 samples/condition). **(H)** Quantification of ELISA for total Tau levels in EV-depleted media from the indicated conditions, with values normalized to CON condition (****P_CON VS. DEX_<0.0001, ***P_DEX VS. DEX + NaClO3_ =0.001, one-way ANOVA with multiple comparisons and Fisher’s LSD test; n=4 samples/condition). (**I-K**) Representative immunoblots (**I**) and quantification **(J-K)** of AT8 and PHF1 immunoreactivity in media from hippocampal neurons treated with vehicle (CON), DEX, or DEX + TDZD-8. Intensity values are normalized to CON condition (***P_CON VS. DEX_ =0.0003, **P_DEX VS. DEX + TDZD_=0.0026 for **J**, ***P_CON VS. DEX_ =0.0001, ***P_DEX VS. DEX + TDZD_ =0.0003 for **K**, one-way ANOVA with multiple comparisons and Fisher’s LSD test; n=3-4 samples/condition). (**L**) Quantification of ELISA for total Tau levels in EV-depleted media from the indicated conditions, with values normalized to CON condition (***P_CON VS. DEX_=0.0003, ****P_DEX VS. DEX + TDZD_ <0.0001, one-way ANOVA with multiple comparisons and Fisher’s LSD test; n=4 samples/condition). (**M-O**) Representative immunoblots (**M**) and quantification **(N-O)** of AT8 and PHF1 immunoreactivity in media from hippocampal neurons treated with vehicle (CON), DEX, or DEX + TTX. Intensity values are normalized to CON condition (****P_CON VS. DEX_ <0.0001, ****P_DEX VS. DEX + TTX_ <0.0001 for **N**, ***P_CON VS. DEX_ =0.0006, **P_DEX VS. DEX + TTX_ =0.0014 for **O**, one-way ANOVA with multiple comparisons and Fisher’s LSD test; n=3-4 samples/condition). (**P**) Quantification of ELISA for total Tau levels in EV-depleted media from the indicated conditions, with values normalized to CON condition (****P_CON VS. DEX_<0.0001, **P_DEX VS. DEX + TTX_ <0.0001, one-way ANOVA with multiple comparisons and Fisher’s LSD test; n=4 samples/condition).

GCs are known to induce Tau hyperphosphorylation via activation of Tau kinases (*e.g.* GSK3β, CDK5)^13, 39–41^ and also to stimulate neuronal firing^42–44^, both of which are reported to enhance Tau secretion^26, 37, 38^. We therefore treated hippocampal neurons with DEX +/- the GSK3β inhibitor TDZD-8 or the Na^+^ channel blocker tetrodotoxin (TTX) to inhibit neuronal firing. Both treatments not only decreased Tau phosphorylation as measured by PHF1 antibody and pS199 Tau ELISA (Fig. **S2J-K**), but also completely blocked the DEX-induced increase in extracellular phospho-and total Tau (Fig. **2I-P**), indicating that GC-mediated Tau secretion is dependent upon Tau phosphorylation and neuronal activity.

Trans-cellular spreading of pathogenic Tau is regarded as a key driver of AD progression^22^. We therefore examined whether Tau secreted in response to high GC levels is internalized by neighboring neurons. For this experiment, media from PS19 ‘donor’ hippocampal neurons treated with vehicle or DEX (1 μM, 48h) was incubated for 48 hours with ‘recipient’ neurons from wild-type mice (Fig. **3A**), and Tau uptake quantified by immunostaining with human-specific anti-Tau13 antibodies. As anticipated based on previous studies^37, 38^, hTau secreted by both control and DEX-treated donor neurons was readily taken up by recipient neurons (Fig. **3B, C**), indicative of its ability to spread trans-cellularly. However, hTau levels were three-fold higher in recipient neurons incubated with medium from DEX-treated donor cells versus vehicle-treated cells (Fig. **3C**). This finding likely reflects increased hTau levels in media following DEX treatment in donor cells, but could also indicate a stimulatory effect of DEX on Tau uptake by recipient cells, or DEX-related toxicity leading to increased membrane permeability to Tau. To investigate these latter possibilities, we treated recipient neurons with DEX for 48 hours during their incubation with (control) donor cell medium and subsequently quantified hTau levels. Interestingly, DEX-treated recipient neurons exhibited similar levels of hTau as their vehicle-treated counterparts (Fig. **3B-C**), and no difference in LDH release (Fig. 3D). These findings indicate that GCs do not stimulate Tau internalization or alter plasma membrane permeability, but rather facilitate Tau spreading by stimulating its secretion.

**Figure 3.**
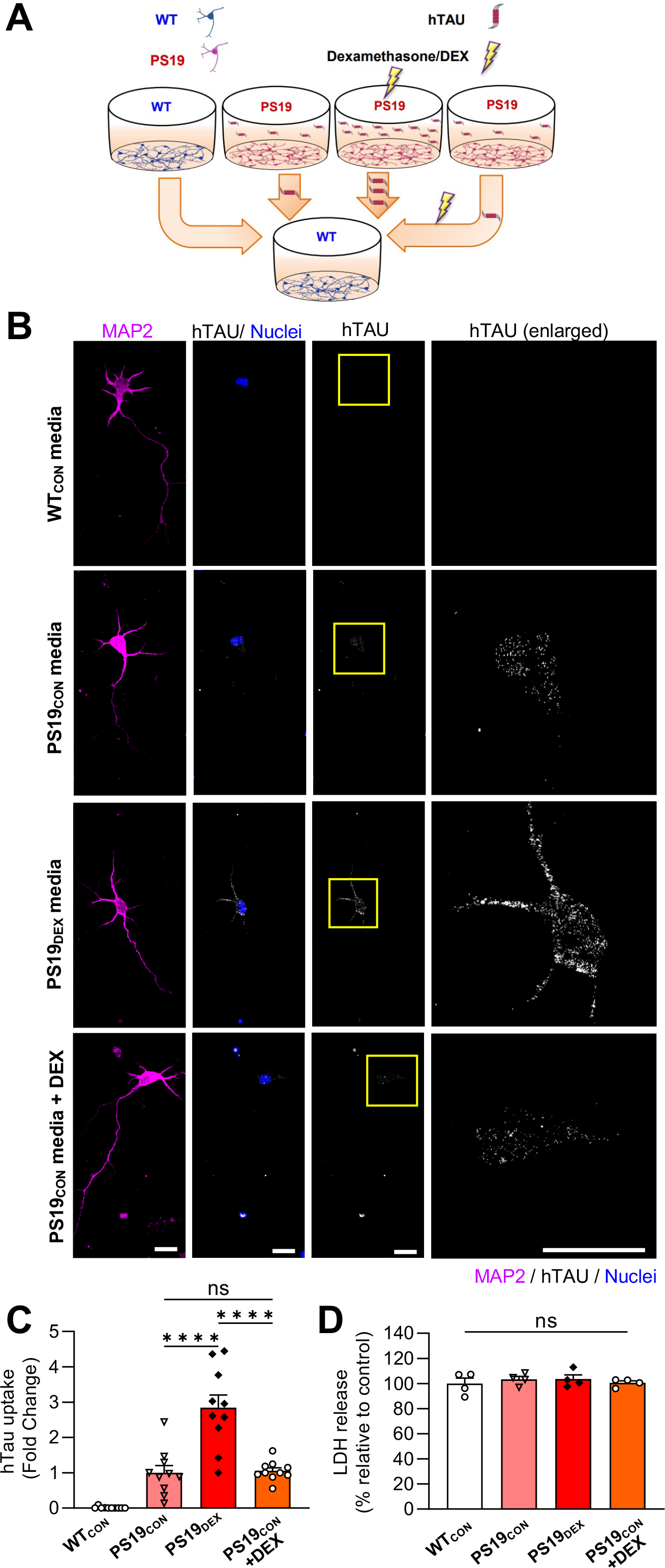
GCs stimulate Tau secretion but not uptake by neurons. (**A**) Schematic diagram of experimental procedure for measuring Tau uptake in cultured neurons. (**B**) Representative images of wild-type recipient hippocampal neurons immunostained for MAP2 (pink) and human-specific Tau (hTau, white), following 48-hour incubation with media from: WT donor neurons treated with vehicle control for 48h (WT_CON_), PS19 donor neurons treated with vehicle for 48h (PS19_CON_), PS19 donor neurons treated with DEX for 48h (PS19_DEX_), or PS19 donor neurons treated with vehicle for 48h, with DEX added to recipient neurons (PS19_CON_ +DEX). Nuclei are stained with DAPI. Scale bar, 25 µm. (**C**) Quantification of Tau uptake by recipient neurons, measured by hTau fluorescence intensity and normalized to PS19_CON_ condition (****P_PS19con VS. PS19dex_ <0.0001, ****P _PS19dex VS. PS19con+DEX_<0.0001; one-way ANOVA with multiple comparisons and Fisher’s LSD test; n=10 fields of view/condition). (**D**) Quantification of LDH release showing no difference between the conditions.

Finally, we evaluated whether high circulating GC levels promote Tau spreading *in vivo*. Here, 4-5 month-old wild-type male mice (3/group) were pre-treated with vehicle, DEX, or DEX + MIF for 7 days, then injected in hippocampal area CA1 (Fig. **4A**) with an adeno-associated virus (AAV) that enables visualization of trans-cellular Tau spreading (AAV.CBA.eGFP.2A.P301L-Tau)^45^. Animals were then treated for an additional 14 days with vehicle (CON), DEX, or DEX + MIF prior to tissue harvest. DEX administration caused a ∼10% loss of body weight during this time period, demonstrating its ability to promote an endocrine response mimicking stress (Fig. **S2L**). After brains from each treatment group were harvested and sectioned, human P301LTau spreading was evaluated by immunostaining with anti-Tau13 antibodies (Fig. **4B, C**). Tau propagation was quantified as in previous studies^45, 46^, by counting hTau^+^/GFP^-^ neurons per mm^2^ near the injection site and calculating the ratio of hTau^+^ cells expressing GFP (GFP/hTau colocalization) (Fig. **4D, E**). Remarkably, the number of hTau^+^/GFP^-^ neurons was dramatically increased in DEX-treated animals compared to CON or DEX + MIF conditions (Fig. **4B, D**), while GFP/hTau colocalization was significantly decreased (Fig. **4B, E**). AAV transduction efficiency (number of GFP^+^ cells per mm^2^) was similar across treatment conditions (Fig. **4F**). Moreover, in DEX-treated animals, hTau was detected in brain areas more than 1000 μm away from GFP^+^ neurons, a phenomenon not observed in the other two groups (Fig. **4C, G**). These data demonstrate that GCs strongly promote Tau secretion and spreading *in vivo*. To assess whether this spreading occurs via type 1 UPS, we initiated a second hTau spreading experiment with epigallocatechin gallate (EGCG), a potent inhibitor of Tau aggregation that attenuates its secretion via type 1 UPS^37, 47^ and can be used *in vivo*, unlike other inhibitors of type 1 UPS^48, 49^. We first verified the ability of EGCG to reduce GC-induced Tau secretion in brain slices (Fig. **S2M-P**). Animals were then subjected to the same experimental paradigm as above, but with EGCG instead of MIF. We again found that DEX provoked a ∼10% loss of body weight, and this phenotype was not rescued by EGCG (similar to MIF treatment; Fig. **S2L**). As predicted by its *ex vivo* efficacy, EGCG administration almost completely prevented DEX-induced Tau spreading in the hippocampus (**Fig. 4H-M**), showing that this process occurs via Tau oligomerization and secretion via type 1 UPS.

**Figure 4.**
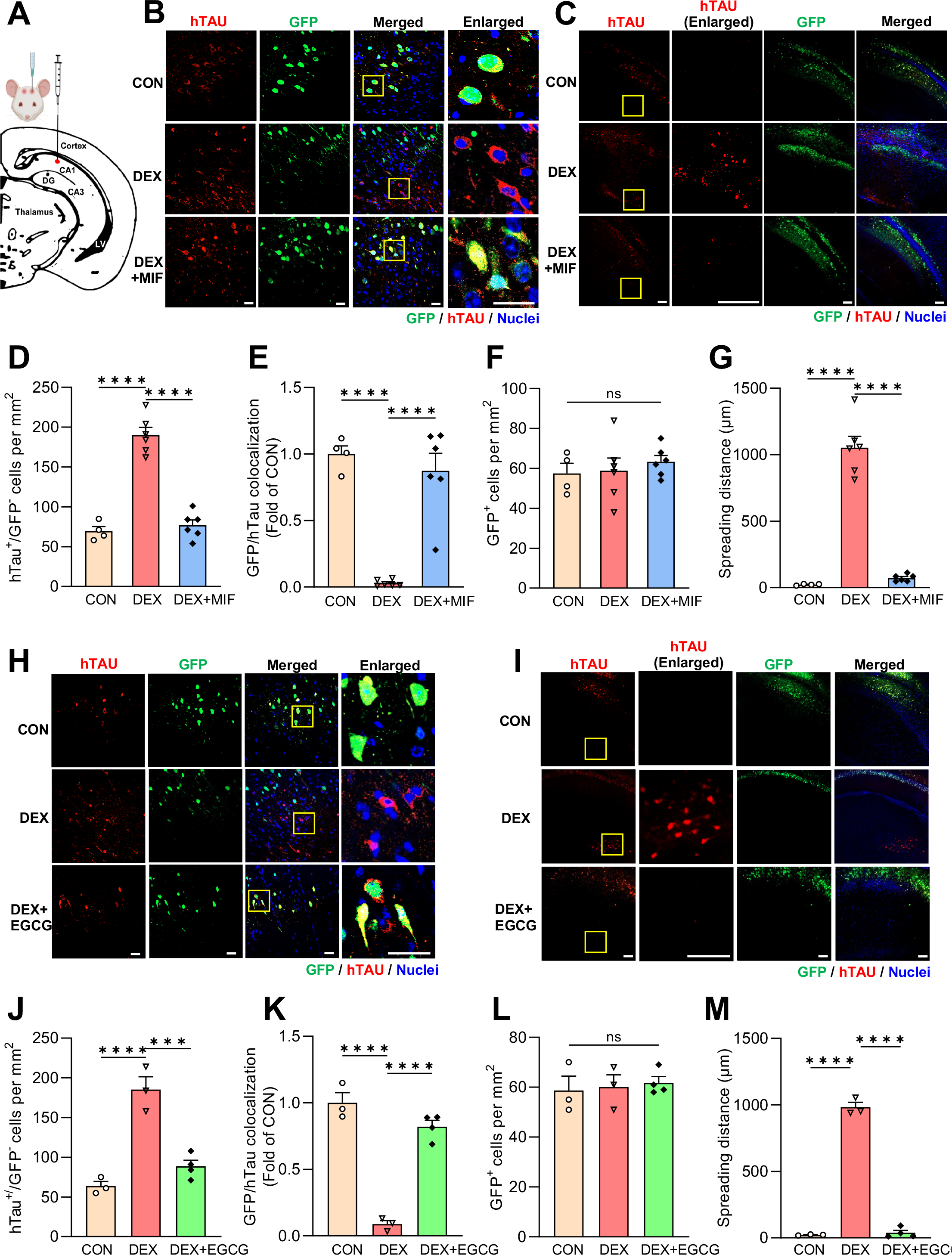
GCs induce Tau spreading *in vivo* through type 1 UPS. (**A**) Schematic diagram indicating the injection site of AAV in murine hippocampal area CA1. (**B**) Representative images showing the colocalization of hTau (red) and GFP (green) in CA1 neurons of mice treated with vehicle (CON), dexamethasone (DEX), or DEX + MIF. Nuclei are stained with DAPI (blue). Right column shows enlarged regions (indicated by yellow boxes). Scale bars, 50 µm. (**C**) Representative images depicting the spreading of hTau (red) from GFP^+^ cells near the injection site in mice treated as indicated. Note hTau spreading beyond the hippocampus in the DEX condition only (yellow box). Scale bars, 200 µm. (**D**) Quantification of hTau^+^/GFP^-^ cells per mm^2^ in mice treated as indicated (\*\*\*\**P*_CON VS. DEX_ <0.0001, \*\*\*\**P*_DEX VS. DEX + MIF_ <0.0001, one-way ANOVA with multiple comparisons and Fisher’s LSD test; *n*=4-6 mice/condition). (**E**) Quantification of the GFP/hTau colocalization ratio in each condition, normalized to CON condition (\*\*\*\**P*_CON VS. DEX_ <0.0001, \*\*\*\**P*_DEX VS. DEX + MIF_ <0.0001, one-way ANOVA with multiple comparisons and Fisher’s LSD test; *n*=4-6 mice/condition). (**F**) Quantification of GFP^+^ cells per mm^2^ in mice treated as indicated (*P*_CON VS. DEX_ =0.8637, *P*_DEX VS. DEX + MIF_ =0.5203, one-way ANOVA with multiple comparisons and Fisher’s LSD test; *n*=4-6 mice/condition). (**G**) Quantification of Tau spreading distance for each condition (\*\*\*\**P*_CON VS. DEX_ <0.0001, \*\*\*\**P*_DEX VS. DEX + MIF_ <0.0001, one-way ANOVA with multiple comparisons and Fisher’s LSD test; *n*=4-6 mice/condition). (**H-I**) Representative images showing hTau/GFP colocalization (**H**) and spreading (**I**) for mice treated as indicated. (**J**) Quantification of hTau^+^/GFP^-^ cells per mm^2^ in mice treated as indicated (\*\*\*\**P*_CON VS. DEX_ <0.0001, \*\*\**P*_DEX VS. DEX + EGCG_=0.0003, one-way ANOVA with multiple comparisons and Fisher’s LSD test; *n*=3-4 mice/condition). (**K**) Quantification of the GFP/hTau colocalization ratio in each condition, normalized to CON condition (\*\*\*\**P*_CON VS. DEX_ <0.0001, \*\*\*\**P*_DEX VS. DEX + EGCG_ <0.0001, one-way ANOVA with multiple comparisons and Fisher’s LSD test; *n*=4 mice/condition). (**L**) Quantification of GFP^+^ cells per mm^2^ in mice treated as indicated (*P*_CON VS. DEX_ =0.8393, *P*_DEX VS. DEX + EGCG_ =0.7763, one-way ANOVA with multiple comparisons and Fisher’s LSD test; *n*=3-4 mice/condition). (**M**) Quantification of Tau spreading distance for each condition (\*\*\*\**P*_CON VS. DEX_ <0.0001, \*\*\*\**P*_DEX VS. DEX + EGCG_ <0.0001, one-way ANOVA with multiple comparisons and Fisher’s LSD test; *n*=3-4 mice/condition).

## Discussion

This work provides the first demonstration that GCs stimulate Tau spreading in the brain, implicating these stress hormones in both the initial stages of Tau pathogenesis, by inducing Tau hyperphosphorylation and aggregation within neurons, and subsequently in the transmission of pathogenic Tau between neurons. We show that GCs stimulate Tau secretion via type 1 UPS, an ATP-independent process requiring interactions between phosphorylated/oligomeric Tau and plasma membrane-associated HSPGs and lipids. Notably, Tau secretion and spreading have also been shown to occur via extracellular vesicles (*i.e.* exosomes and ectosomes) and to be mediated by other brain cell types including microglia^24, 50^. Additional work will be required to determine whether GCs also stimulate Tau propagation via these mechanisms.

An intriguing finding of this study is that EGCG, a catechin found at high levels in green tea leaves, blocks GC-induced Tau spreading *in vivo*. EGCG is an inhibitor of Tau oligomerization and aggregation as well as its secretion via type 1 UPS, suggesting that this is an important mode of GC-driven Tau propagation. However, EGCG also alters lipid membrane properties^51, 52^ and could alter Tau secretion/uptake via this mechanism. Other drugs that block type 1 UPS, such as NaClO_3_ and methyl-β-cyclodextrin, have similarly pleiotropic effects (and further cannot be used *in vivo* due to their blood brain barrier impermeability and toxicity, respectively^48, 49^), making it difficult to definitively demonstrate that secretion via type 1 UPS is the primary driver of GC-induced Tau propagation *in vivo*. However, given the relative amount of Tau reported to undergo secretion in vesicle-free form (∼90%)^22^, we think it is reasonable to assume that type 1 UPS contributes substantially to GC-induced Tau spreading *in vivo*.

Our experiments further reveal that GCs promote Tau secretion by stimulating GSK3β- mediated Tau phosphorylation. GSK3β, a brain-enriched serine/threonine kinase implicated in Tau pathogenesis in AD, phosphorylates multiple Tau residues, including those detected by our ELISA (S199) and immunoblotting (Ser202/Thr205, S396/S404) assays^53, 54^. Indeed, we observe that GCs selectively increase secretion of pS199 Tau compared to total Tau, and the GSK3β inhibitor TDZD-8 effectively blocks GC-mediated Tau secretion in hippocampal neurons. Interestingly, we also find that neuronal activity is critical for this process, as treatment with TTX to inhibit action potential firing prevents GC-induced Tau phosphorylation as well as its secretion. These data are in line with other studies reporting that neuronal activity, in the form of depolarization or NMDA receptor activation, stimulates Tau phosphorylation^55, 56^. On the other hand, phosphorylated Tau can also exert effects on neuronal activity. In particular, the mislocalization of phospho-Tau species to dendritic spines in response to stress/GCs has been suggested to induce aberrant neuronal firing/excitotoxicity via Fyn kinase-mediated opening of NMDA receptors, leading to Ca^2+^ influx^14, 15, 56^. These findings suggest the existence of a positive feedback loop, wherein GC-induced neuronal activity promotes Tau phosphorylation, which in turn induces the synaptic mistargeting of phospho-Tau species that stimulate additional neuronal activity to continue this cycle. However, since GCs are known to activate multiple Tau kinases, including CDK5 and GSK3β^13, 39–41^, and also to stimulate the firing of cortical and hippocampal glutamatergic neurons on a rapid timescale (1-4h after application)^42–44^, it may be challenging to fully disentangle the causality of these events.

Cumulatively, our data show that GC-mediated phosphorylation and oligomerization of Tau stimulates its vesicle-free secretion and trans-cellular spreading via type 1 UPS. While questions remain about how Tau phosphorylation is precipitated by GCs, and how stress/GCs impact other forms of Tau propagation in the brain, this work provides some of the first mechanistic insight into how high GC levels accelerate pathogenic Tau spreading in AD and other tauopathies.

## Acknowledgements

This work was supported by NIH grants R01NS080967 to C.L.W. and RF1AG069941 to C.L.W. and I.S., and Portuguese Foundation for Science & Technology PhD fellowship PD/BD/135271/2017 to P.G. We would like to thank Dr. Carol Troy and members of the Troy lab for guidance in the use of their mouse stereotactic frame.

## Author Contributions

Q.Y., F.D., and C.L.W. designed the research; Q.Y., F.D., I.B., and P.G. performed experiments and analyzed data; I.S. and C.L.W. supervised experiments; C.L.W. wrote the manuscript; Q.Y. and F.D. prepared figures; Q.Y., F.D., P.G., I.S., and C.L.W. edited the manuscript. All authors read and approved the final manuscript.

## STAR METHODS

## KEY RESOURCES TABLE

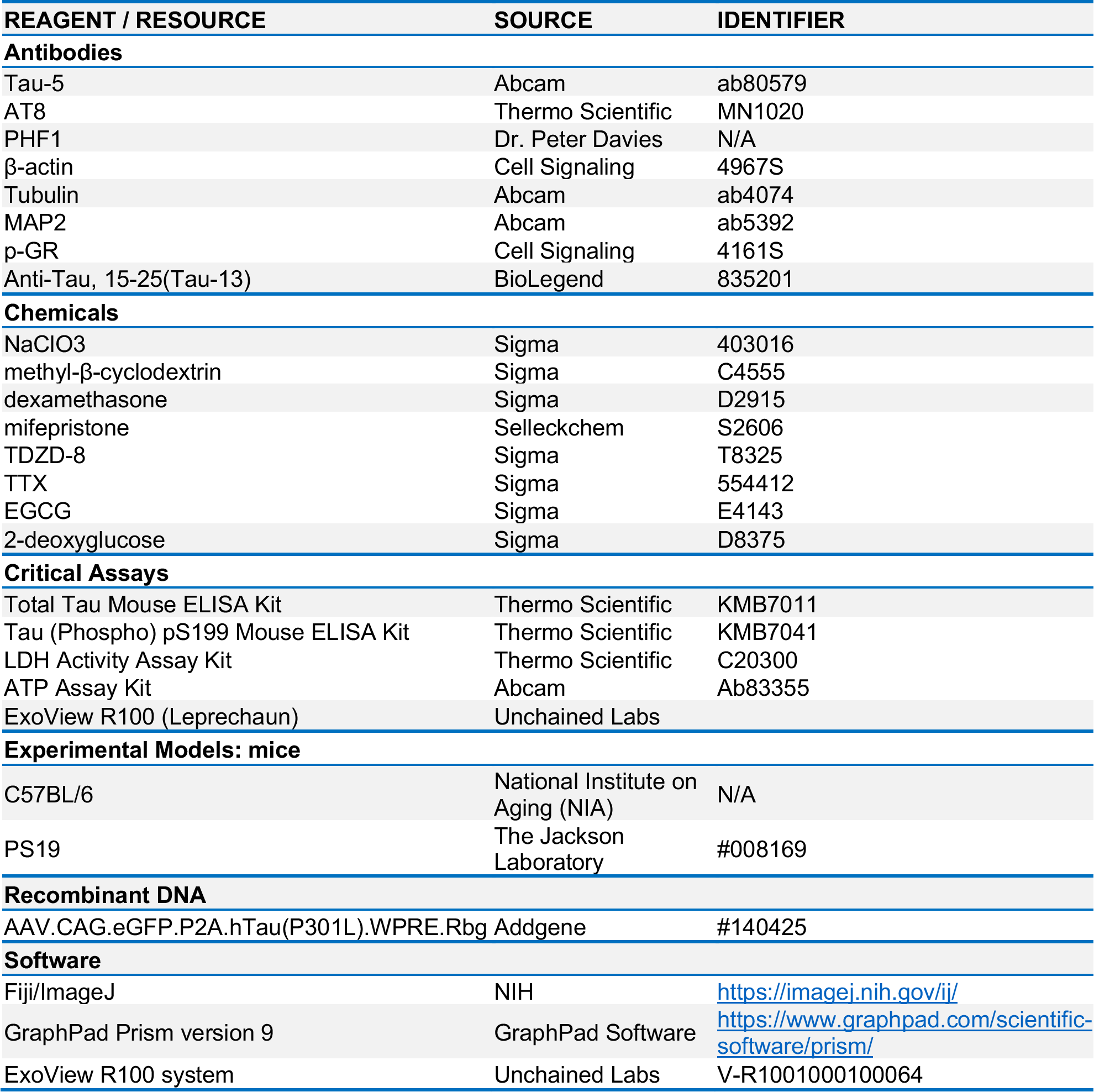

## CONTACT FOR REAGENT AND RESOURCE SHARING

Further information and requests for resources and reagents should be directed to and will be fulfilled by the Lead Contact, Clarissa Waites

## EXPERIMENTAL MODEL AND SUBJECT DETAILS

### Animals

Male and female C57BL/6 mice (obtained from the National Institute on Aging) and PS19 mice (obtained from The Jackson Laboratory; strain #008169) between the ages of 4-5 months were maintained under standard laboratory conditions with ad libitum access to food and water. All animal studies were carried out with the approval of the Columbia Institutional Animal Care and Use Committee (IACUC) in accordance with the National Institutes of Health guidelines for animal care.

## METHOD DETAILS

### Primary hippocampal culture

Primary mouse hippocampal neurons were prepared from postnatal day 0 wild-type or PS19 mice, as described previously^57^, and maintained in 24-well plates with Neurobasal medium supplemented with B27, 600 μM L-glutamine, and antibiotic-antimycotic (all from ThermoFisher/Life Technologies). At 11-12 days *in vitro* (DIV), media was replaced with new media containing 0.5% B27 supplement and treated as follows: control (50% PEG400 diluted into media (vehicle for dex/mifepristone), dexamethasone (dex, 1µM) for 48 hours, mifepristone (5 µM) for 1-hour pre-treatment + 48 hours together with dex, NaClO_3_ (50mM) for 24-hour pre-treatment + 48 hours together with dex, methyl-β-cyclodextrin (1mM) for 24-hour pre-treatment + 48 hours with dex. For all conditions, media was collected at 14 DIV. For LDH measurements, media was collected from 14 DIV PS19 neurons with indicated treatments.

### Brain slice perfusion

Brains were harvested from mice sacrificed via cervical dislocation without anesthesia followed by decapitation^58^. Brain slices including cortex and hippocampus (coronal sections; 400μm) were cut and maintained in an interface chamber at 29°C and perfused with artificial cerebrospinal fluid (ACSF) continuously bubbled with 95% O_2_ and 5% CO_2_. ACSF composition was as follows: 124 mM NaCl, 4.4 mM KCl, 1 mM Na_2_HPO_4_, 25 mM NaHCO_3_, 2 mM CaCl_2_, 2 mM MgCl_2_ and 10 mM glucose. ACSF was collected from *ex vivo* brain slices after the following treatments: control (50% PEG400 diluted into ACSF (dex/mifepristone vehicle)), dexamethasone (5 µM) for 4 hours, mifepristone (5 µM) for 1-hour pre-treatment + 4 hours with dex, NaClO_3_ (100mM), methyl-β- cyclodextrin (2.5mM), or EGCG (50µM) for 1.5-hour pre-treatment + 4 hours with dex.

### Media/ACSF preparation for immunoblot and ELISA

When indicated, the cell culture media or ACSF were centrifuged for 20min at 2000g to eliminate cell debris, then concentrated using Pierce™ Protein Concentrators PES with 30K molecular-weight cutoff (Thermo Scientific, 88531). To deplete extracellular vesicle (EVs), media/ACSF was subjected to sequential centrifugation steps: 30 min at 10,000 g, 30 min at 21,000 g, and finally 70 min at 100,000 g. The remaining supernatant was used for immunoblotting and ELISA.

### CSF collection

Five-month-old C57BL/6 mice (13/group; 10 male and 3 female) were administered dexamethasone (D2915, Sigma; 5mg/kg per day, dissolved in PBS, by intraperitoneal/i.p. injection), and mifepristone/RU486 (S2606, Selleckchem; 10mg/kg per day, dissolved in 50% PEG400 in PBS, by i.p.) for 15 days. Control animals received injections of 50% PEG400 diluted in PBS. Following this treatment regimen, mice were euthanized by isoflurane and CSF was collected from the cisterna magna using a glass capillary.

### Chronic unpredictable stress, brain tissue collection, and media harvest

Three-to four-month-old wild-type animals (C57BL/6J) were housed in groups of 5–6 per cage under standard environmental conditions with ad libitum access to food and water. For the chronic unpredictable stress (CUS) protocol, animals were subjected to different stressors (i.e. 3 hours overcrowding, 3 hours rocking platform, 3 hours restraint, 30 min hairdryer; one stressor per day) that were chosen randomly to prevent habituation, over a period of six weeks. Following the CUS protocol, animals were euthanized, brain tissue was immediately macrodissected and incubated in EV-release medium (Neurobasal medium, 1% Glutamax, 1% Anti-anti; ThermoFisher) for 16h at 37°C, 5% CO_2_. Five hemi-cortices were pooled to obtain each cortical sample while hippocampi from 5 mouse brains were pooled into each hippocampal sample. After the incubation period, media was collected and subject to extracellular vesicle depletion as described above (Media/ACSF preparation).

### ExoView Imager Analysis

The characterization and quantification of exosomes in hippocampal culture media were performed according to the manufacturer’s instructions^59^. Briefly, chips containing capture probes coated with antibodies against two exosome-enriched tetraspanins, CD81 and CD9, were pre-scanned to acquire baseline particle adhesion prior to sample incubation. Media samples were diluted to fall within the dynamic range of the Exoview R100 instrument (Unchained Labs), and incubated overnight at room temperature on the pre-scanned chips in a sealed 24-well plate. The chips were then washed to remove any non-captured material, incubated for 1 hour at room temperature with fluorescently-conjugated antibodies against CD9, CD63, and CD81, washed again, dried, and then scanned with the ExoView R100 system to obtain data on particle counts, size, and exosome surface membrane protein profiles. For each capture probe (CD9 and CD81), background particle readout is subtracted from the final particle count to produce a final exosome count readout.

### Immunoblotting

The concentrated media/ACSF with extracellular vesicle (EV) depletion were prepared in 4x Laemmli buffer and then boiled for 5 min, followed by SDS/PAGE (10% Tris-Glycine gel; XP00105BOX, Invitrogen), then transferred to a nitrocellulose membrane (10600001, Amersham). After blocking in TBST buffer (20 mM Tris-HCl, 150 mM sodium chloride, 0.1% Tween-20) containing 5% (wt/vol) nonfat dry milk for 1 h at room temperature, the membrane was incubated with primary antibodies overnight at 4°C, then with secondary antibodies for 1 h at room temperature. The following antibodies were used: Tau5 (ab80579, Abcam), AT8: anti-phospho-Tau pSer202/Thr205 (MN1020, ThermoFisher Scientific), PHF-1: anti-phospho-Tau pSer396/Ser404 Tau (from Dr. Peter Davies), β-actin (4967S, cell signaling), anti-Tubulin (ab4074, Abcam). IRDye 800CW goat anti-mouse IgG secondary antibody (P/N: 926-32210, LI-COR), IRDye 680CW goat anti-rabbit IgG secondary antibody (P/N: 926-68071, LI-COR). Membranes were visualized by Odyssey Infrared Imager (model 9120, LI-COR Biosciences), and relative optical densities of bands determined by Fiji/ImageJ software.

### ELISA

EV-depleted media/ACSF samples (50 µL volume) were used for measurement of Tau concentration by a mouse-specific total Tau ELISA kit (KMB7011, Thermo Scientific) or pS199 Tau ELISA kit (KMB7041) according to manufacturer’s instructions.

### Tau uptake assay

Media was collected from donor WT or PS19 neurons treated with vehicle control or dex (1µM) for 48 hours. The media from these cultures was then depleted of EVs as described above and transferred to naïve recipient wild-type neurons for a 48-hour incubation. For one condition, recipient neurons were also treated with dex (1µM) during this time. Following incubation, recipient cells were washed three times with cold 1x PBS and fixed with 4% paraformaldehyde as previously described^57^. The uptake of hTau was then detected by immunostaining with MAP2 and Tau13 antibodies as described below.

### Immunofluorescence staining of brain slices, cultured neurons

Floating brain sections or fixed primary neurons were immunostained as previously described ^57^. Briefly, fixed neurons or slices cut at 35 μm on a vibratome (VT1000S; Leica) were incubated overnight with the following primary antibodies: mouse Anti-Tau, 15-25(Tau-13) antibody (1:1000, 835201, BioLegend) and chicken MAP2 (1:5000, ab5392, Abcam). They were then incubated for 1 h with secondary antibodies (Alexa Fluor® 594 anti-mouse IgG, and Alexa Fluor® 633 anti-chicken IgG, 1:2000 dilution). Coverslips were mounted with VectaShield (Vector Laboratories) and sealed with clear nail polish. Images were acquired with a 63X objective (Neofluar, NA 1.4) on a Zeiss LSM 800 confocal microscope running Zen2 software. The images were manually measured and quantified using the auto-threshold settings in Fiji/ImageJ software.

### AAV injection procedure

The AAV.CBA.eGFP.2A.P301L-Tau plasmid, a gift from Bradley Hyman (Addgene plasmid #140425; http://n2t.net/addgene:140425;RRID:Addgene_140425), was packaged into AAV8 serotype by University of Pennsylvania Viral Vector Core. Prior to AAV injection, male/female mice (3-4/group) were administered dex (5 mg/kg, i.p.injection) +/- mifepristone (10 mg/kg, i.p. injection) or dex (5 mg/kg, i.p. injection) +/- EGCG (20 mg/kg, i.p. injection) for 7 days. Stereotactic AAV injections were performed under standard aseptic surgery conditions as previously described^46^. Briefly, mice were anaesthetized with isoflurane (2%), placed in a stereotactic frame (digital stereotaxic device, Stoelting Co.), and injected bilaterally with 2 μl of AAV in hippocampal region CA1 (at the following coordinates relative to Bregma: A/P −2.7 mm, M/L ±2 mm, D/V −1.5 mm) with a 10 μl Hamilton syringe at a rate of 0.25 μl/min by a Nano-injector system (Stoelting microsyringe pump, Stoelting Co.). The needle was kept in place for an additional 5 min. Afterwards, the skin over the injection site was sutured and mice were placed on a warming pad during their recovery from anesthesia. Mice were then administered dex with or without mifepristone or EGCG for an additional 14 days prior to euthanasia and brain harvest. Control animals received daily i.p. injections of 50% PEG400 in PBS (dex/mifepristone vehicle) or PBS (dex/EGCG vehicle).

### Quantification and statistical analysis

All values were expressed as the mean ± SEM. All graphing and statistical analyses were performed using GraphPad Prism (GraphPad Prism9.Ink). Statistical details of experiments are provided in the figure legends. Statistical analyses were performed with unpaired, two-tailed t-test or one-way ANOVA, with appropriate corrections for unequal variances and multiple comparisons. A minimum of 3 independent replicates were used for all experiments. Values of *p* < 0.05 were considered statistically significant. *p<0.05, **p<0.01, ***p<0.001,****p<0.0001.

## Supplementary Material

## Supplemental Figure Legends

**Figure S1.**
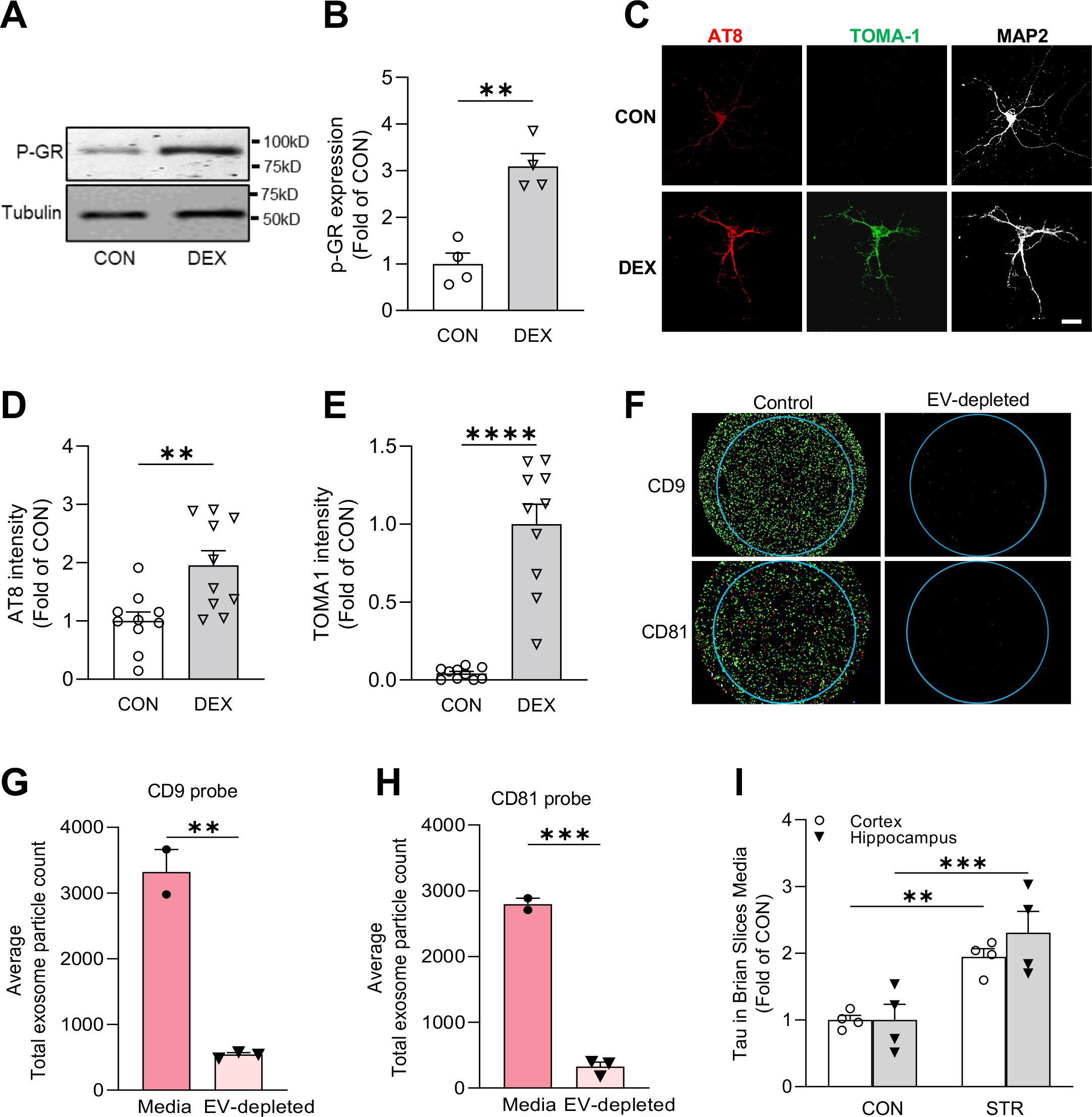
Additional control experiments. (**A-B**) Immunoblots and quantification of phospho-GR in hippocampal lysates treated as indicated. Intensity is normalized to the CON condition (**P_CON VS. DEX_ =0.0011, unpaired two-tailed t-test, n=4 samples/condition). (**C-E**) Representative images (**C**) and quantification (**D-E**) of AT8 (red) and TOMA-1 (green) fluorescence intensity in MAP2 positive (grey) cultured hippocampal neurons treated with as indicated. Scale bars, 25 µm. Intensity values are normalized to CON condition (**P _CON VS. DEX_ =0.0042 for **D**, ****P_CON VS. DEX_ <0.0001 for **E**, unpaired two-tailed t-test, n=10 fields of view/condition). (**F**) Images of EVs on CD9- and CD81-coated capture probes, from hippocampal culture media +/- EV depletion. (**G-H**) Quantification of particle counts on CD9 (**G**) and CD81 (**H**) capture probes from the indicated conditions (**P_media VS. EV depleted_=0.0017 for **G**, ***P_media VS. EV depleted_=0.0002 for **H**, unpaired, two-tailed t-test; n=2-3 samples/condition). (**I**) Quantification of ELISA for total Tau levels from media containing cortical or hippocampal brain slices from mice subjected to chronic unpredictable stress (STR) compared to control (CON) conditions (\*\**P*_CON VS. STR_ =0.0079, \*\*\**P*_CON VS. STR_ =0.0009, unpaired, two-tailed t-test; n=4 samples/condition).

**Figure S2.**
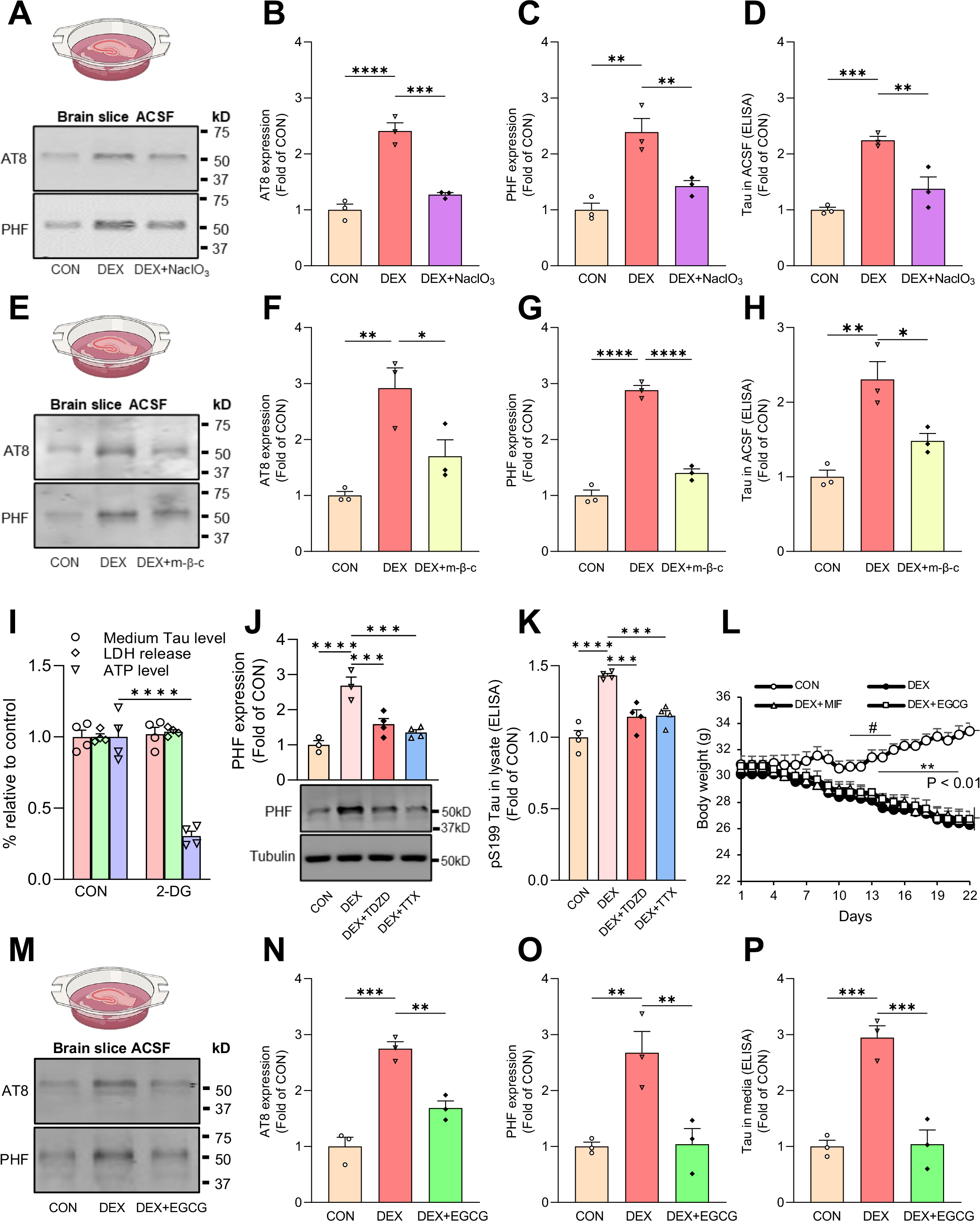
Tau secretion from *ex vivo* brain slices. (**A-C**) Representative immunoblots (**A**) and quantification **(B-C)** of AT8 and PHF1 immunoreactivity in ACSF from brain slices perfused with vehicle (CON), DEX, or DEX + NaClO_3_. Intensity values are normalized to CON condition (****P _CON VS. DEX_ <0.0001, ***P_DEX VS. DEX + NaClO3_ =0.0003 for **B**, **P_CON VS. DEX_ =0.0011, **P_DEX VS. DEX + NaClO3_ =0.0064 for **C,** one-way ANOVA with multiple comparisons and Fisher’s LSD test; n=3 samples/condition). **(D)** Quantification of ELISA for total Tau levels in EV-depleted ACSF from the indicated conditions, with values normalized to CON condition (***P_CON VS. DEX_=0.0005, **P_DEX VS. DEX + NaClO3_ =0.0035, one-way ANOVA with multiple comparisons and Fisher’s LSD test; n=3 samples/condition). (**E-G**) Representative immunoblots (**E**) and quantification **(F-G)** of AT8 and PHF1 immunoreactivity in ACSF from brain slices perfused with vehicle (CON), DEX, or DEX + methyl-b-cyclodextrin (m-β-c). Intensity values are normalized to CON condition (**P _CON VS. DEX_ =0.0025, **P_DEX VS. DEX + m-β-c_ =0.0196 for **F**, ****P_CON VS. DEX_ <0.0001, ****P_DEX VS. DEX + m-β-c_ <0.0001 for **G**,one-way ANOVA with multiple comparisons and Fisher’s LSD test; n=3 samples/condition). **(H)** Quantification of ELISA for total Tau levels in EV-depleted ACSF from the indicated conditions, with values normalized to CON condition (**P_CON VS. DEX_=0.0011, *P_DEX VS. DEX + NaClO3_ =0.0102, one-way ANOVA with multiple comparisons and Fisher’s LSD test; n=3 samples/condition). (**I**) Quantification of total Tau (ELISA), LDH, and ATP levels in EV-depleted media from primary neurons treated as indicated, with values normalized to CON condition (P_CON VS. 2-DG_=0.7746 for Tau ELISA assay, P_CON VS. 2-DG_=0.5949 for LDH assay, ****P_CON VS. 2-DG_<0.0001 for ATP assay, unpaired, two-tailed t-test; n=4 samples/condition). (**J**) Representative immunoblots and quantification of PHF1 Tau in lysates from hippocampal neurons treated as indicated (****P _CON VS. DEX_ <0.0001, ***P_DEX VS. DEX + TDZD_ =0.0006, ***P_DEX VS. DEX + TTX_ =0.0001, one-way ANOVA with multiple comparisons and Fisher’s LSD test; n=3-4 samples/condition). (**K**) Quantification of ELISA for pS199 Tau levels in lysate with the indicated conditions, with values normalized to CON condition (****P _CON VS. DEX_ <0.0001, ***P_DEX VS. DEX + TDZD_ =0.0002, ***P_DEX VS. DEX + TTX_ =0.0002, one-way ANOVA with multiple comparisons and Fisher’s LSD test; n=4 samples/condition). (**L**) Quantification of body weight loss in 4-5-month-old male/female mice treated for 21 days with CON, DEX, DEX + MIF, or DEX + EGCG [P _CON VS. DEX, DEX+MIF, DEX+EGCG_<0.001 (days 21), **P _CON VS. DEX_ <0.01 (days 15-20), ^#^P _CON VS. DEX+MIF, DEX+EGCG_<0.05 (days 12-14), 2-way ANOVA with multiple comparisons and Fisher’s LSD test; n=3-7 mice/condition). (**M-O**) Representative immunoblots (**M**) and quantification **(N-O)** of AT8 and PHF1 immunoreactivity in ACSF from brain slices perfused with vehicle (CON), DEX, or DEX + EGCG. Intensity values are normalized to CON condition (***P _CON VS. DEX_ =0.0001, **P_DEX VS. DEX + EGCG_ =0.0018 for **N**, ****P_CON VS. DEX_ =0.0052, ****P_DEX VS. DEX + EGCG_=0.0059 for **O**, one-way ANOVA with multiple comparisons and Fisher’s LSD test; n=3 samples/condition). **(P)** Quantification of ELISA for total Tau levels in EV-depleted ACSF from the indicated conditions, with values normalized to CON condition (***P_CON VS. DEX_=0.0005, *P_DEX VS. DEX + EGCG_ =0.0006, one-way ANOVA with multiple comparisons and Fisher’s LSD test; n=3 samples/condition).

## References

1. Caruso, A., Nicoletti, F., Gaetano, A., and Scaccianoce, S. (2019). Risk Factors for Alzheimer’s Disease: Focus on Stress. Front Pharmacol 10, 976. 10.3389/fphar.2019.00976.

2. Mravec, B., Horvathova, L., and Padova, A. (2018). Brain Under Stress and Alzheimer’s Disease. Cell Mol Neurobiol 38, 73–84. 10.1007/s10571-017-0521-1.

3. Machado, A., Herrera, A.J., de Pablos, R.M., Espinosa-Oliva, A.M., Sarmiento, M., Ayala, A., Venero, J.L., Santiago, M., Villaran, R.F., Delgado-Cortes, M.J., et al. (2014). Chronic stress as a risk factor for Alzheimer’s disease. Rev Neurosci 25, 785–804. 10.1515/revneuro-2014-0035.

4. Johansson, L., Guo, X., Waern, M., Ostling, S., Gustafson, D., Bengtsson, C., and Skoog, I. (2010). Midlife psychological stress and risk of dementia: a 35-year longitudinal population study. Brain 133, 2217–2224. 10.1093/brain/awq116.

5. Huang, C.W., Lui, C.C., Chang, W.N., Lu, C.H., Wang, Y.L., and Chang, C.C. (2009). Elevated basal cortisol level predicts lower hippocampal volume and cognitive decline in Alzheimer’s disease. J Clin Neurosci 16, 1283–1286. 10.1016/j.jocn.2008.12.026.

6. Mejia, S., Giraldo, M., Pineda, D., Ardila, A., and Lopera, F. (2003). Nongenetic factors as modifiers of the age of onset of familial Alzheimer’s disease. International psychogeriatrics 15, 337–349.

7. Green, K.N., Billings, L.M., Roozendaal, B., McGaugh, J.L., and LaFerla, F.M. (2006). Glucocorticoids increase amyloid-beta and tau pathology in a mouse model of Alzheimer’s disease. J Neurosci 26, 9047–9056. 10.1523/JNEUROSCI.2797-06.2006.

8. Han, B., Yu, L., Geng, Y., Shen, L., Wang, H., Wang, Y., Wang, J., and Wang, M. (2016). Chronic Stress Aggravates Cognitive Impairment and Suppresses Insulin Associated Signaling Pathway in APP/PS1 Mice. J Alzheimers Dis 53, 1539–1552. 10.3233/JAD-160189.

9. Jeong, Y.H., Park, C.H., Yoo, J., Shin, K.Y., Ahn, S.M., Kim, H.S., Lee, S.H., Emson, P.C., and Suh, Y.H. (2006). Chronic stress accelerates learning and memory impairments and increases amyloid deposition in APPV717I-CT100 transgenic mice, an Alzheimer’s disease model. FASEB J 20, 729–731. 10.1096/fj.05-4265fje.

10. Carroll, J.C., Iba, M., Bangasser, D.A., Valentino, R.J., James, M.J., Brunden, K.R., Lee, V.M., and Trojanowski, J.Q. (2011). Chronic stress exacerbates tau pathology, neurodegeneration, and cognitive performance through a corticotropin-releasing factor receptor-dependent mechanism in a transgenic mouse model of tauopathy. J Neurosci 31, 14436–14449. 10.1523/JNEUROSCI.3836-11.2011.

11. Baglietto-Vargas, D., Chen, Y., Suh, D., Ager, R.R., Rodriguez-Ortiz, C.J., Medeiros, R., Myczek, K., Green, K.N., Baram, T.Z., and LaFerla, F.M. (2015). Short-term modern life-like stress exacerbates Abeta-pathology and synapse loss in 3xTg-AD mice. J Neurochem 134, 915–926. 10.1111/jnc.13195.

12. Dong, H., Goico, B., Martin, M., Csernansky, C.A., Bertchume, A., and Csernansky, J.G. (2004). Modulation of hippocampal cell proliferation, memory, and amyloid plaque deposition in APPsw (Tg2576) mutant mice by isolation stress. Neuroscience 127, 601–609. 10.1016/j.neuroscience.2004.05.040.

13. Sotiropoulos, I., Catania, C., Pinto, L.G., Silva, R., Pollerberg, G.E., Takashima, A., Sousa, N., and Almeida, O.F. (2011). Stress acts cumulatively to precipitate Alzheimer’s disease-like tau pathology and cognitive deficits. J Neurosci 31, 7840–7847. 10.1523/JNEUROSCI.0730-11.2011.

14. Pinheiro, S., Silva, J., Mota, C., Vaz-Silva, J., Veloso, A., Pinto, V., Sousa, N., Cerqueira, J., and Sotiropoulos, I. (2015). Tau Mislocation in Glucocorticoid-Triggered Hippocampal Pathology. Mol Neurobiol. 10.1007/s12035-015-9356-2.

15. Lopes, S., Vaz-Silva, J., Pinto, V., Dalla, C., Kokras, N., Bedenk, B., Mack, N., Czisch, M., Almeida, O.F., Sousa, N., and Sotiropoulos, I. (2016). Tau protein is essential for stress-induced brain pathology. Proc Natl Acad Sci U S A 113, E3755–3763. 10.1073/pnas.1600953113.

16. Roberson, E.D., Scearce-Levie, K., Palop, J.J., Yan, F., Cheng, I.H., Wu, T., Gerstein, H., Yu, G.Q., and Mucke, L. (2007). Reducing endogenous tau ameliorates amyloid beta-induced deficits in an Alzheimer’s disease mouse model. Science 316, 750–754. 10.1126/science.1141736.

17. Vossel, K.A., Zhang, K., Brodbeck, J., Daub, A.C., Sharma, P., Finkbeiner, S., Cui, B., and Mucke, L. (2010). Tau reduction prevents Abeta-induced defects in axonal transport. Science 330, 198. 10.1126/science.1194653.

18. Braak, H., and Braak, E. (1991). Neuropathological stageing of Alzheimer-related changes. Acta Neuropathol 82, 239–259.

19. Bejanin, A., Schonhaut, D.R., La Joie, R., Kramer, J.H., Baker, S.L., Sosa, N., Ayakta, N., Cantwell, A., Janabi, M., Lauriola, M., et al. (2017). Tau pathology and neurodegeneration contribute to cognitive impairment in Alzheimer’s disease. Brain 140, 3286–3300. 10.1093/brain/awx243.

20. Nelson, P.T., Alafuzoff, I., Bigio, E.H., Bouras, C., Braak, H., Cairns, N.J., Castellani, R.J., Crain, B.J., Davies, P., Del Tredici, K., et al. (2012). Correlation of Alzheimer disease neuropathologic changes with cognitive status: a review of the literature. J Neuropathol Exp Neurol 71, 362–381. 10.1097/NEN.0b013e31825018f7.

21. DeLeo, A.M., and Ikezu, T. (2017). Extracellular Vesicle Biology in Alzheimer’s Disease and Related Tauopathy. J Neuroimmune Pharmacol. 10.1007/s11481-017-9768-z.

22. Brunello, C.A., Merezhko, M., Uronen, R.L., and Huttunen, H.J. (2020). Mechanisms of secretion and spreading of pathological tau protein. Cell Mol Life Sci 77, 1721–1744. 10.1007/s00018-019-03349-1.

23. Dujardin, S., Begard, S., Caillierez, R., Lachaud, C., Delattre, L., Carrier, S., Loyens, A., Galas, M.C., Bousset, L., Melki, R., et al. (2014). Ectosomes: a new mechanism for non-exosomal secretion of tau protein. PLoS One 9, e100760. 10.1371/journal.pone.0100760.

24. Asai, H., Ikezu, S., Tsunoda, S., Medalla, M., Luebke, J., Haydar, T., Wolozin, B., Butovsky, O., Kugler, S., and Ikezu, T. (2015). Depletion of microglia and inhibition of exosome synthesis halt tau propagation. Nat Neurosci 18, 1584–1593. 10.1038/nn.4132.

25. Wang, Y., Balaji, V., Kaniyappan, S., Kruger, L., Irsen, S., Tepper, K., Chandupatla, R., Maetzler, W., Schneider, A., Mandelkow, E., and Mandelkow, E.M. (2017). The release and trans-synaptic transmission of Tau via exosomes. Molecular neurodegeneration 12, 5. 10.1186/s13024-016-0143-y.

26. Pooler, A.M., Phillips, E.C., Lau, D.H., Noble, W., and Hanger, D.P. (2013). Physiological release of endogenous tau is stimulated by neuronal activity. EMBO Rep 14, 389–394. 10.1038/embor.2013.15.

27. Chai, X., Dage, J.L., and Citron, M. (2012). Constitutive secretion of tau protein by an unconventional mechanism. Neurobiol Dis 48, 356–366. 10.1016/j.nbd.2012.05.021.

28. Fontaine, S.N., Zheng, D., Sabbagh, J.J., Martin, M.D., Chaput, D., Darling, A., Trotter, J.H., Stothert, A.R., Nordhues, B.A., Lussier, A., et al. (2016). DnaJ/Hsc70 chaperone complexes control the extracellular release of neurodegenerative-associated proteins. EMBO J 35, 1537–1549. 10.15252/embj.201593489.

29. Wu, J.W., Hussaini, S.A., Bastille, I.M., Rodriguez, G.A., Mrejeru, A., Rilett, K., Sanders, D.W., Cook, C., Fu, H., Boonen, R.A., et al. (2016). Neuronal activity enhances tau propagation and tau pathology in vivo. Nat Neurosci 19, 1085–1092. 10.1038/nn.4328.

30. Guix, F.X., Corbett, G.T., Cha, D.J., Mustapic, M., Liu, W., Mengel, D., Chen, Z., Aikawa, E., Young-Pearse, T., Kapogiannis, D., et al. (2018). Detection of Aggregation-

31. Competent Tau in Neuron-Derived Extracellular Vesicles. Int J Mol Sci 19.10.3390/ijms19030663.

31. Wegmann, S., Nicholls, S., Takeda, S., Fan, Z., and Hyman, B.T. (2016). Formation, release, and internalization of stable tau oligomers in cells. J Neurochem 139, 1163–1174. 10.1111/jnc.13866.

32. Faure, J., Lachenal, G., Court, M., Hirrlinger, J., Chatellard-Causse, C., Blot, B., Grange, J., Schoehn, G., Goldberg, Y., Boyer, V., et al. (2006). Exosomes are released by cultured cortical neurones. Mol Cell Neurosci 31, 642–648. 10.1016/j.mcn.2005.12.003.

33. Lopes, S., Teplytska, L., Vaz-Silva, J., Dioli, C., Trindade, R., Morais, M., Webhofer, C., Maccarrone, G., Almeida, O.F.X., Turck, C.W., et al. (2017). Tau Deletion Prevents Stress-Induced Dendritic Atrophy in Prefrontal Cortex: Role of Synaptic Mitochondria. Cereb Cortex 27, 2580–2591. 10.1093/cercor/bhw057.

34. Du, F., Yu, Q., Swerdlow, R.H., and Waites, C.L. (2023). Glucocorticoid-driven mitochondrial damage stimulates Tau pathology. Brain. 10.1093/brain/awad127.

35. Thery, C., Amigorena, S., Raposo, G., and Clayton, A. (2006). Isolation and characterization of exosomes from cell culture supernatants and biological fluids. Current protocols in cell biology / editorial board, Juan S. Bonifacino … [et al.] Chapter 3, Unit 3 22. 10.1002/0471143030.cb0322s30.

36. Keerthikumar, S., Gangoda, L., Liem, M., Fonseka, P., Atukorala, I., Ozcitti, C., Mechler, A., Adda, C.G., Ang, C.S., and Mathivanan, S. (2015). Proteogenomic analysis reveals exosomes are more oncogenic than ectosomes. Oncotarget 6, 15375–15396. 10.18632/oncotarget.3801.

37. Merezhko, M., Brunello, C.A., Yan, X., Vihinen, H., Jokitalo, E., Uronen, R.L., and Huttunen, H.J. (2018). Secretion of Tau via an Unconventional Non-vesicular Mechanism. Cell reports 25, 2027–2035 e2024. 10.1016/j.celrep.2018.10.078.

38. Katsinelos, T., Zeitler, M., Dimou, E., Karakatsani, A., Muller, H.M., Nachman, E., Steringer, J.P., Ruiz de Almodovar, C., Nickel, W., and Jahn, T.R. (2018). Unconventional Secretion Mediates the Trans-cellular Spreading of Tau. Cell reports 23, 2039–2055. 10.1016/j.celrep.2018.04.056.

39. Sotiropoulos, I., Catania, C., Riedemann, T., Fry, J.P., Breen, K.C., Michaelidis, T.M., and Almeida, O.F. (2008). Glucocorticoids trigger Alzheimer disease-like pathobiochemistry in rat neuronal cells expressing human tau. J Neurochem 107, 385–397. 10.1111/j.1471-4159.2008.05613.x.

40. Yi, J.H., Brown, C., Whitehead, G., Piers, T., Lee, Y.S., Perez, C.M., Regan, P., Whitcomb, D.J., and Cho, K. (2017). Glucocorticoids activate a synapse weakening pathway culminating in tau phosphorylation in the hippocampus. Pharmacol Res 121, 42–51. 10.1016/j.phrs.2017.04.015.

41. Dey, A., Hao, S., Wosiski-Kuhn, M., and Stranahan, A.M. (2017). Glucocorticoid-mediated activation of GSK3beta promotes tau phosphorylation and impairs memory in type 2 diabetes. Neurobiol Aging 57, 75–83. 10.1016/j.neurobiolaging.2017.05.010.

42. Beck, S.G., List, T.J., and Choi, K.C. (1994). Long-and short-term administration of corticosterone alters CA1 hippocampal neuronal properties. Neuroendocrinology 60, 261–272. 10.1159/000126758.

43. Joels, M. (2009). Stress, the hippocampus, and epilepsy. Epilepsia 50, 586–597. 10.1111/j.1528-1167.2008.01902.x.

44. Krugers, H.J., Alfarez, D.N., Karst, H., Parashkouhi, K., van Gemert, N., and Joels, M. (2005). Corticosterone shifts different forms of synaptic potentiation in opposite directions. Hippocampus 15, 697–703. 10.1002/hipo.20092.

45. Wegmann, S., Bennett, R.E., Delorme, L., Robbins, A.B., Hu, M., McKenzie, D., Kirk, M.J., Schiantarelli, J., Tunio, N., Amaral, A.C., et al. (2019). Experimental evidence for the age dependence of tau protein spread in the brain. Sci Adv 5, eaaw6404. 10.1126/sciadv.aaw6404.

46. Rauch, J.N., Luna, G., Guzman, E., Audouard, M., Challis, C., Sibih, Y.E., Leshuk, C., Hernandez, I., Wegmann, S., Hyman, B.T., et al. (2020). LRP1 is a master regulator of tau uptake and spread. Nature 580, 381–385. 10.1038/s41586-020-2156-5.

47. Wobst, H.J., Sharma, A., Diamond, M.I., Wanker, E.E., and Bieschke, J. (2015). The green tea polyphenol (-)-epigallocatechin gallate prevents the aggregation of tau protein into toxic oligomers at substoichiometric ratios. FEBS Lett 589, 77–83. 10.1016/j.febslet.2014.11.026.

48. Gosselet, F., Loiola, R.A., Roig, A., Rosell, A., and Culot, M. (2021). Central nervous system delivery of molecules across the blood-brain barrier. Neurochem Int 144, 104952. 10.1016/j.neuint.2020.104952.

49. Ali, S.N., Arif, A., Ansari, F.A., and Mahmood, R. (2022). Cytoprotective effect of taurine against sodium chlorate-induced oxidative damage in human red blood cells: an ex vivo study. Amino Acids 54, 33–46. 10.1007/s00726-021-03121-5.

50. Merezhko, M., Uronen, R.L., and Huttunen, H.J. (2020). The Cell Biology of Tau Secretion. Front Mol Neurosci 13, 569818. 10.3389/fnmol.2020.569818.

51. Patra, S.K., Rizzi, F., Silva, A., Rugina, D.O., and Bettuzzi, S. (2008). Molecular targets of (-)-epigallocatechin-3-gallate (EGCG): specificity and interaction with membrane lipid rafts. J Physiol Pharmacol 59 *Suppl 9*, 217–235.

52. Sun, Y., Hung, W.C., Chen, F.Y., Lee, C.C., and Huang, H.W. (2009). Interaction of tea catechin (-)-epigallocatechin gallate with lipid bilayers. Biophys J 96, 1026–1035. 10.1016/j.bpj.2008.11.007.

53. Balasubramaniam, M., Mainali, N., Bowroju, S.K., Atluri, P., Penthala, N.R., Ayyadevera, S., Crooks, P.A., and Shmookler Reis, R.J. (2020). Structural modeling of GSK3beta implicates the inactive (DFG-out) conformation as the target bound by TDZD analogs. Scientific reports 10, 18326. 10.1038/s41598-020-75020-w.

54. Sayas, C.L., and Avila, J. (2021). GSK-3 and Tau: A Key Duet in Alzheimer’s Disease. Cells 10. 10.3390/cells10040721.

55. Pierrot, N., Santos, S.F., Feyt, C., Morel, M., Brion, J.P., and Octave, J.N. (2006). Calcium-mediated transient phosphorylation of tau and amyloid precursor protein followed by intraneuronal amyloid-beta accumulation. J Biol Chem 281, 39907–39914. 10.1074/jbc.M606015200.

56. Hu, Z., Ondrejcak, T., Yu, P., Zhang, Y., Yang, Y., Klyubin, I., Kennelly, S.P., Rowan, M.J., and Hu, N.W. (2023). Do tau-synaptic long-term depression interactions in the hippocampus play a pivotal role in the progression of Alzheimer’s disease? Neural Regen Res 18, 1213–1219. 10.4103/1673-5374.360166.

57. Du, F., Yu, Q., Yan, S., Hu, G., Lue, L.F., Walker, D.G., Wu, L., Yan, S.F., Tieu, K., and Yan, S.S. (2017). PINK1 signalling rescues amyloid pathology and mitochondrial dysfunction in Alzheimer’s disease. Brain 140, 3233–3251. 10.1093/brain/awx258.

58. Yu, Q., Wang, Y., Du, F., Yan, S., Hu, G., Origlia, N., Rutigliano, G., Sun, Q., Yu, H., Ainge, J., et al. (2018). Overexpression of endophilin A1 exacerbates synaptic alterations in a mouse model of Alzheimer’s disease. Nat Commun 9, 2968. 10.1038/s41467-018-04389-0.

59. Breitwieser, K., Koch, L.F., Tertel, T., Proestler, E., Burgers, L.D., Lipps, C., Adjaye, J., Furst, R., Giebel, B., and Saul, M.J. (2022). Detailed Characterization of Small Extracellular Vesicles from Different Cell Types Based on Tetraspanin Composition by ExoView R100 Platform. Int J Mol Sci 23. 10.3390/ijms23158544.

